# ARMOR: an <u>A</u>utomated <u>R</u>eproducible <u>MO</u>dular workflow for preprocessing and differential analysis of <u>R</u>NA-seq data

**DOI:** 10.1101/575951

**Authors:** Stephany Orjuela, Ruizhu Huang, Katharina M. Hembach, Mark D. Robinson, Charlotte Soneson

## Abstract

The extensive generation of RNA sequencing (RNA-seq) data in the last decade has resulted in a myriad of specialized software for its analysis. Each software module typically targets a specific step within the analysis pipeline, making it necessary to join several of them to get a single cohesive workflow. Multiple software programs automating this procedure have been proposed, but often lack modularity, transparency or flexibility. We present ARMOR, which performs an end-to-end RNA-seq data analysis, from raw read files, via quality checks, alignment and quantification, to differential expression testing, geneset analysis and browser-based exploration of the data. ARMOR is implemented using the Snakemake workflow management system and leverages conda environments; Bioconductor objects are generated to facilitate downstream analysis, ensuring seamless integration with many R packages. The workflow is easily implemented by cloning the GitHub repository, replacing the supplied input and reference files and editing a configuration file. Although we have selected the tools currently included in ARMOR, the setup is modular and alternative tools can be easily integrated.

## 1 Introduction

Since the first high-throughput RNA-seq experiments about a decade ago, there has been a tremendous development in the understanding of the characteristic features of the collected data, as well as the methods used for the analysis. Today there are vetted, well-established algorithms and tools available for many aspects of RNA-seq data analysis (Conesa et al. 2016; Van Den Berge et al. 2018). In this study, we capitalize on this knowledge and present a modular, light-weight RNA-seq workflow covering the most common parts of a typical end-to-end RNA-seq data analysis. The application is implemented using the Snakemake workflow management system (Köster and Rahmann 2012), and allows the user to easily perform quality assessment, adapter trimming, genome alignment, transcript and gene abundance quantification, differential expression analysis and geneset analyses with a simple command, after specifying the required reference files and information about the experimental design in a configuration file. Reproducibility is ensured via the use of conda environments, and all relevant log files are retained for transparency. The output is provided in state-of-the-art R/Bioconductor classes, ensuring interoperability with a broad range of Bioconductor packages. In particular, we provide a template to facilitate browser-based interactive visualization of the quantified abundances and the results of the statistical analyses with iSEE (Rue-Albrecht et al. 2018).

Among already existing pipelines for automated reference-based RNA-seq analysis, several focus either on the preprocessing and quality control steps (He et al. 2018; Ewels et al. 2018; Tsyganov et al. 2018), or on the downstream analysis and visualization of differentially expressed genes (Marini 2018; Monier et al. 2019; Powell 2018), or do not provide a single framework for the preprocessing and downstream analysis (Steinbaugh et al. 2018). Some workflows are based on predefined reference files and can only quantify abundances for human or mouse (Torre, Lachmann, and Ma’ayan 2018; Cornwell et al. 2018; Wang 2018). Additionally, workflows that conduct differential gene expression analysis often do not allow comparisons between more than two groups, or more complex experimental designs (Girke 2018; Cornwell et al. 2018). Some existing pipelines only provide a graphical user interface to design and execute fully automated analyses (Hung et al. 2018; Guerler et al. 2018). In addition to reference-based tools, there are also pipelines that perform *de novo* transcriptome assembly before downstream analysis (e.g. https://dib-lab.github.io/eelpond/).

ARMOR performs both preprocessing and downstream statistical analysis of the RNA-seq data, building on standard statistical analysis methods and commonly used data containers. It distinguishes itself from existing workflows in several ways: (i) Its modularity, reflected in its fully and easily customizable framework. (ii) The transparency of the output and analysis, meaning that all code is accessible and can be modified by the user. (iii) The seamless integration with downstream analysis and visualization packages, especially those within Bioconductor (Huber et al. 2015). (iv) The ability to specify any fixed-effect experimental design and any number of contrasts, in a standardized format. (v) The inclusion of a test for differential transcript usage in addition to differential gene expression analysis. While high-performance computing environments and cloud computing are not specifically targeted, Snakemake enables the usage of a cluster without the need to modify the workflow itself.

In general, we do not advocate fully automated analysis. All rigorous data analyses need exploratory steps and spot checks at various steps throughout the process, to ensure that data is of sufficient quality and to spot potential errors (e.g., sample mislabelings). ARMOR handles the automation of "bookkeeping" tasks, such as running the correct sequence of software for all samples, and compiling the data and reports in standardized formats. If errors are identified, the workflow can re-run only the parts that need to be updated.

ARMOR is available from https://github.com/csoneson/ARMOR.

## 2 Materials and Methods

### 2.1 Overview

The ARMOR workflow is designed to perform an end-to-end analysis of bulk RNA-seq data, starting from FASTQ files with raw sequencing reads (Figure 1). Reads first undergo quality control with FastQC (https://www.bioinformatics.babraham.ac.uk/projects/fastqc/) and (optionally) adapter trimming using TrimGalore! (https://www.bioinformatics.babraham.ac.uk/projects/trim_galore/), before being mapped to a transcriptome index using Salmon (Patro et al. 2017) and (optionally) aligned to the genome using STAR (Dobin et al. 2013). Estimated transcript abundances from Salmon are imported into R using the tximeta package (Soneson, Love, and Robinson 2015; Love et al. 2019) and analyzed for differential gene expression and (optionally) differential transcript usage with edgeR (Robinson, McCarthy, and Smyth 2010) and DRIMSeq (Nowicka and Robinson 2016). The quantifications, provided metadata, and results from the statistical analyses are exported as SingleCellExperiment objects (Lun and Risso 2019) ensuring interoperability with a large part of the Bioconductor ecosystem (Huber et al. 2015). Quantification and QC results are summarized in a MultiQC report (Ewels et al. 2016). Other tools can be easily exchanged for those listed above by modifying the Snakefile and/or the template analysis code.

**Figure 1:**
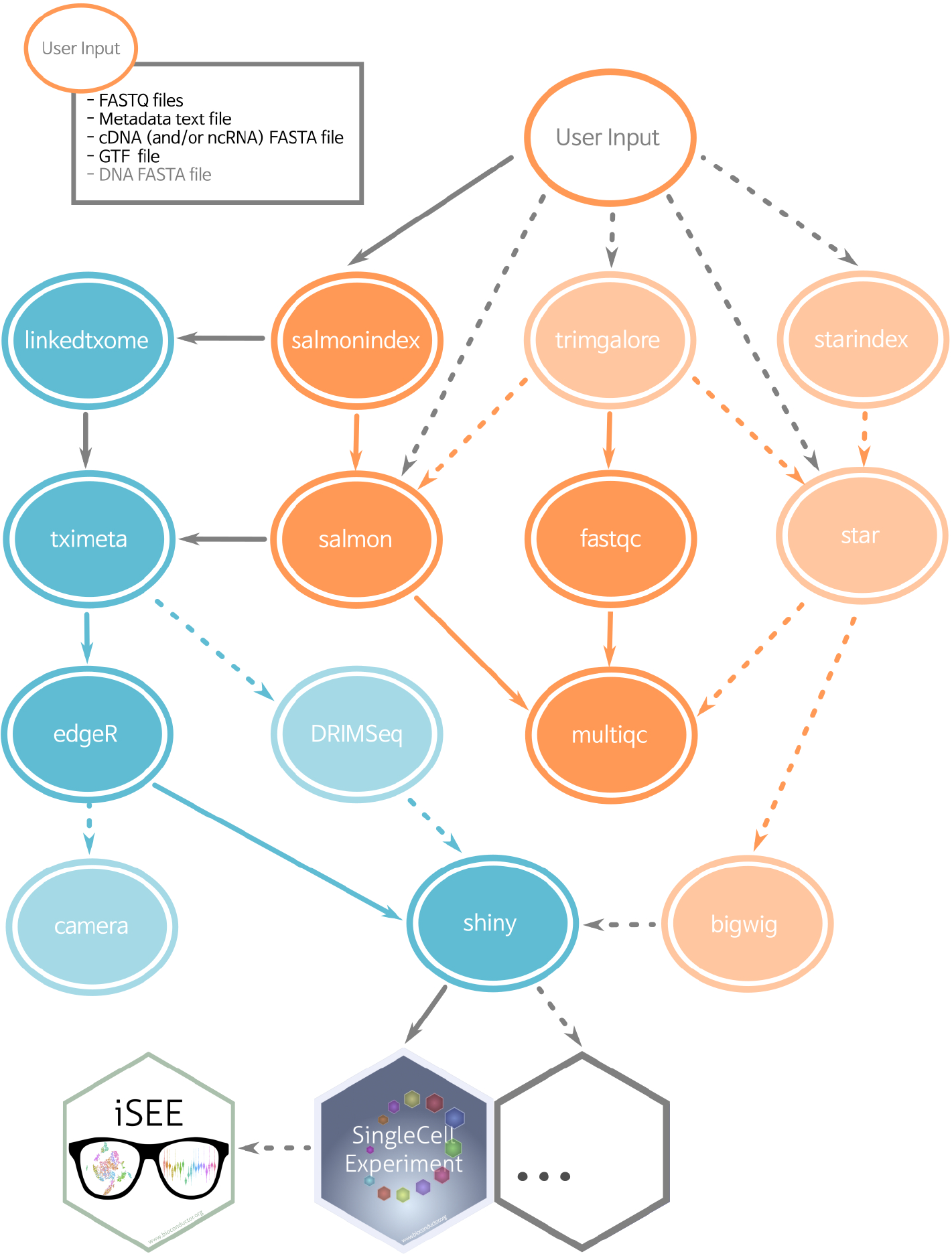
Simplified directed acyclic graph (DAG) of the ARMOR workflow. Blue ellipses are rules run in R, orange ellipses from software called as shell commands. Dashed lines and light-colored ellipses are optional rules, controlled in the configuration file. By default all rules are executed.

### 2.2 Input file specification

ARMOR can be used to analyze RNA-seq data from any organism for which a reference transcriptome, and (optionally) an annotated reference genome are available from either Ensembl (Zerbino et al. 2018) or GENCODE (Frankish et al. 2019). Paths to the reference files, as well as the FASTQ files with the sequencing reads, are specified by the user in a configuration file. In addition, the user prepares a metadata file, a tab-delimited text file listing the name of the samples, the library type (single-or paired-end) and any other covariates that will be used for the statistical analysis. The `checkinputs` rule in the `Snakefile` can be executed to make sure all the input files and the parameters in the configuration file have been correctly specified.

### 2.3 Workflow execution

ARMOR is implemented as a modular Snakemake (Köster and Rahmann 2012) workflow, and the execution of the individual steps is controlled by the provided `Snakefile`. Snakemake will automatically keep track of the dependencies between the different parts of the workflow; rerunning the workflow will thus only regenerate results that are out of date or missing given these dependencies. Via a set of variables specified in the configuration file, the user can easily decide to include or exclude the optional parts of the workflow (shaded ellipses in Figure 1). By adding or modifying targets in the `Snakefile`, users can include any additional or specialized types of analyses that are not covered by the original workflow.

By default, all software packages that are needed for the analysis will be installed in an auto-generated conda (Grüning et al. 2018) environment, which will be automatically activated by Snakemake before the execution of each rule. The desired software versions can be specified in the provided environment file. If the user prefers, local installations of (all or a subset of) the required software can also be used (as described in Software management).

### 2.4 Software management

First, the user needs to have a recent version of snakemake and conda installed. There is a range of possibilities to manage the software for the ARMOR workflow. The recommended option is to allow conda and the workflow itself to manage everything, including the installation of the needed R packages. The workflow is executed this way with the command

~~~
snakemake --use-conda
~~~

The first time the workflow is run, the conda environments will be generated and all necessary software will be installed. Any subsequent invocations of the workflow from this directory will use these generated environments. An alternative option is to use ARMOR’s envs/environment.yaml file to create a conda environment that can be manually activated, by running the command

~~~
conda env create --name ARMOR \
--file envs/environment.yaml
conda activate ARMOR
~~~

The second command activates the environment. Once the environment is activated, ARMOR can be run by simply typing

~~~
snakemake
~~~

Additionally, the user can circumvent the use of conda, and make sure that all software is already available and in the user’s `PATH`. For this, Snakemake and the tools listed in `envs/environment.yaml` need to be manually installed, in addition to a recent version of R and the R packages listed in `scripts/install_pkgs.R`.

For either of the options mentioned above, the `useCondaR` flag in the configuration file controls whether a local R installation, or a conda installed R, will be used. If `useCondaR` is set to False, the path to a local R installation (e.g., `Rbin:<path>`) must be specified in the configuration file, along with the path to the R package library (e.g., `R_LIBS_USER="<path>"`) in the .Renviron file. If the specified R library does not contain the required packages, Snakemake will try to install them (i.e., write permissions would be needed). ARMOR has been tested on macOS and Linux systems.

### 2.5 Statistical analysis

ARMOR uses the quasi-likelihood framework of edgeR (Robinson, McCarthy, and Smyth 2010; Lun, Chen, and Smyth 2016) to perform tests for differential gene expression, camera (Wu and Smyth 2012) to perform associated geneset analysis, and DRIMSeq (Nowicka and Robinson 2016) to test for differential transcript usage between conditions. All code to perform the statistical analyses is provided in Rmark-down templates (Allaire et al. 2018; Xie, Allaire, and Grolemund 2018), which are executed at runtime. This setup gives the user flexibility to use any experimental design supported by these tools, and to test any contrast(s) of interest, by specifying these in the configuration file using standard R syntax, e.g.,

~~~
design:"~ 0 + group"
contrast:groupA-groupB
~~~

Arbitrarily complex designs and multiple contrasts are supported. In addition, by editing the template code, users can easily configure the analysis, add additional plots, or even replace the statistical test if desired. After compilation, all code used for the statistical analysis, together with the results and version information for all packages used, is retained in a standalone html report, ensuring transparency and reproducibility and facilitating communication of the results.

### 2.6 Output files

The output files from all steps in the ARMOR workflow are stored in a user-specified output directory, together with log files for each step, including relevant software version information. The results from the statistical analyses are combined with the transcript- and gene-level quantifications and saved as SingleCellExperiment objects (Lun and Risso 2019), ensuring easy integration with a large number of Bioconductor packages for downstream analysis and visualization. For example, the results can be interactively explored using the iSEE package (Rue-Albrecht et al. 2018) and a template is provided for this.

### 2.7 Multiple project management

When managing multiple projects, the user might run ARMOR in multiple physical locations (i.e., clone the repository in separate places). `snakemake –-use-conda` will create a separate conda environment in each subdirectory, which means the installed software may be duplicated. If space is a concern, building and activating a conda environment (with conda `env create`) may be beneficial. It is also possible to explicitly specify the path to the desired `config.yaml` configuration file when `snakemake` is called:

~~~
snakemake --configfile config.yaml
~~~

In this way a separate `config.yaml` file can be kept for each project, which may be useful when running multiple projects.

By taking advantage of the Snakemake framework, ARMOR makes file and software organization relatively autonomous. Although we recommend using a file structure similar to the one used for the example data provided in the repository (Figure 2), and managing all the software for a project in a conda environment, the user is free to use the same environment for different datasets, even if the files are located in several folders. This alternative is more of a "software-based" structure than the "project-based" structure we present with the pipeline. Either structure has its advantages and depending on the use case and level of expertise, both can be easily implemented using ARMOR.

**Figure 2:**
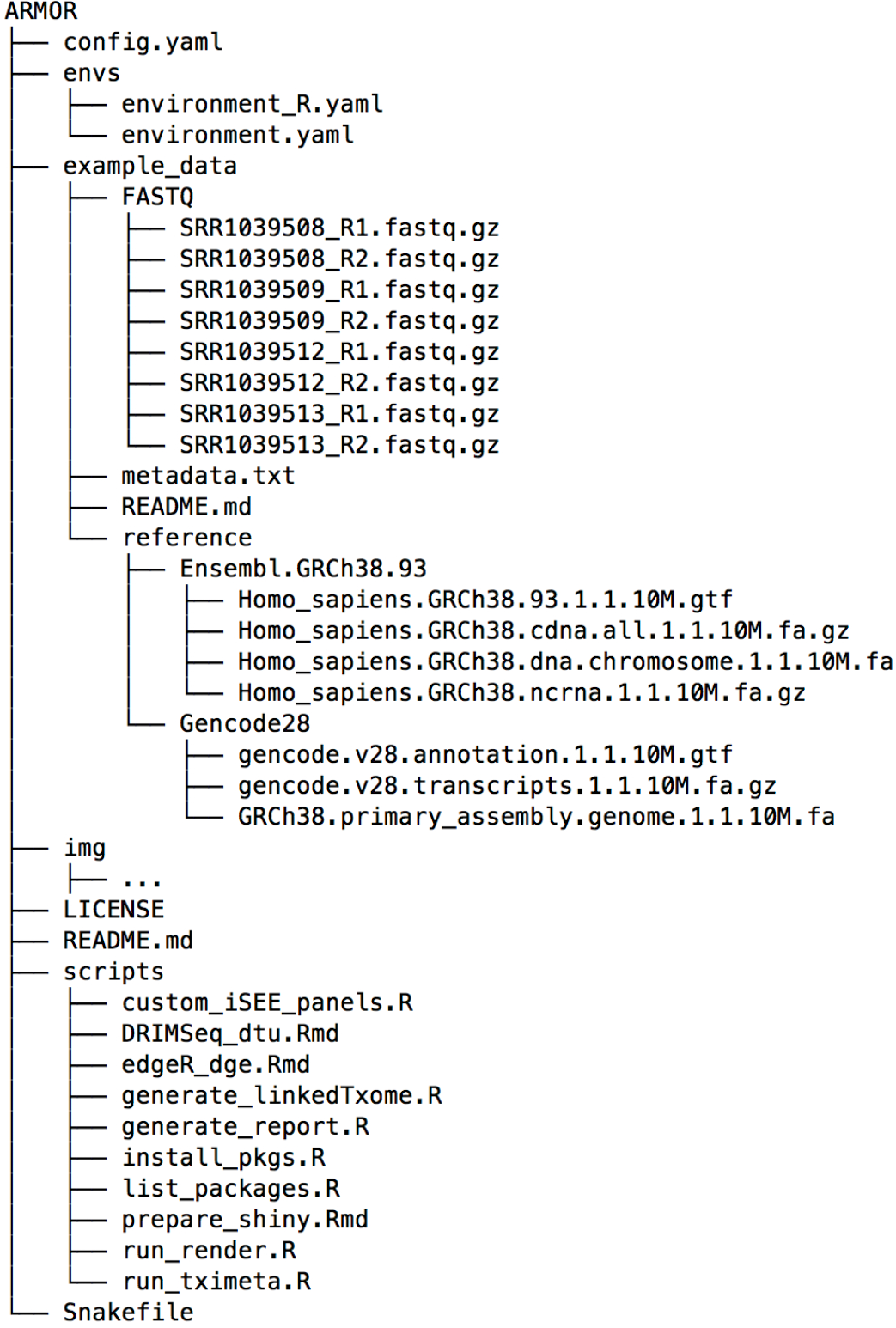
The files and directory structure that make up the ARMOR workflow.

### 2.8 Code availability

ARMOR is available (under MIT license) from https://github.com/csoneson/ARMOR, which also contains a wiki (https://github.com/csoneson/ARMOR/wiki) detailing the usage of the workflow, and a detailed walk-through of an example analysis.

## 3 Results and Discussion

### 3.1 The ARMOR skeleton

Figure 2 shows the set of files contained within the ARMOR workflow, and what is downloaded to the user’s computer when the repository is cloned.

The `example_data` directory represents a (runnable) template of a very small dataset, which is useful for testing the software setup and the system as well as for having a structure to copy for a real project. The provided `config.yaml` file is pre-configured for this example dataset. We recommend that users prepare their own `config.yaml` and a similar directory structure to `example_data`, with the raw FASTQ files and reference sequence and annotation information in subfolders, perhaps using symbolic links if such files are already available in another location. We present an independent example below in the Real data walk-through section.

Once everything is set up, running `snakemake`, which operates on the rules in the `Snakefile`, will construct the hierarchy of instructions to execute, given the specifications in the config.yaml file. Snake-make automatically determines the dependencies between the rules and will invoke the instructions in a logical order. The scripts and envs directories, and the `Snakefile` itself, should not need to be modified, unless the user wants to customize certain aspects of the pipeline.

### 3.2 Real data walk-through

Here, we illustrate the practical usage of ARMOR on a bulk RNA-seq dataset from a study on Wnt signalling (Doumpas et al. 2019). For each of three genetic backgrounds (HEK 293T, dBcat and d4TCF), there are six samples: three biological replicates each of untreated cells, and cells after Wnt pathway stimulation (using the GSK3 inhibitor CHIRON99021). An R script (download_files.R, which can be found at https://github.com/csoneson/ARMOR/blob/chiron_realdataworkflow/E-MTAB-7029/download_files.R) was written to download the FASTQ files with raw reads from ArrayExpress (https://www.ebi.ac.uk/arrayexpress/experiments/E-MTAB-7029/), and create a metadata table detailing the type of library and experimental condition for each sample (Table 1). This table was saved as a tab-delimited text file named `metadata.txt`.

**Table 1:** Metadata table for the Wnt signalling data

| names | type | condition |
| --- | --- | --- |
| Q10-Chir-1_R1 | SE | d4Tcf__chir |
| Q10-Chir-2_R1 | SE | d4Tcf__chir |
| Q10-Chir-3_R1 | SE | d4Tcf__chir |
| b-cat-KO-Chir-1_R1 | SE | dBcat__chir |
| b-cat-KO-Chir-2_R1 | SE | dBcat__chir |
| b-cat-KO-Chir-3_R1 | SE | dBcat__chir |
| WT-Chir-1_R1 | SE | WT__chir |
| WT-Chir-2_R1 | SE | WT__chir |
| WT-Chir-3_R1 | SE | WT__chir |
| Q10-unstim-1_R1 | SE | d4Tcf__unstim |
| Q10-unstim-2_R1 | SE | d4Tcf__unstim |
| Q10-unstim-3_R1 | SE | d4Tcf__unstim |
| b-cat-KO-ustim-1_R1 | SE | dBcat__unstim |
| b-cat-KO-ustim-2_R1 | SE | dBcat__unstim |
| b-cat-KO-ustim-3_R1 | SE | dBcat__unstim |
| WT-unstim-1_R1 | SE | WT__unstim |
| WT-unstim-2_R1 | SE | WT__unstim |
| WT-unstim-3_R1 | SE | WT__unstim |

The raw data and reference files were organized into a directory, E-MTAB-7029, with the structure according to Figure 3. The default config.yaml downloaded with the workflow was copied into a new file called `config_E-MTAB-7029.yaml` and edited to reflect the location of these files. In addition, the read length was set and the experimental design was specified as " ~ 0 + condition", where the condition information will be taken from metadata.txt. Then, a set of contrasts of interest (e.g., conditiond4Tcf chir-conditiond4Tcf unstim) were specified, as well as the set of genesets to use. The final configuration file can be viewed at https://github.com/csoneson/ARMOR/blob/chiron_realdataworkflow/config_E-MTAB-7029.yaml.

**Figure 3:**
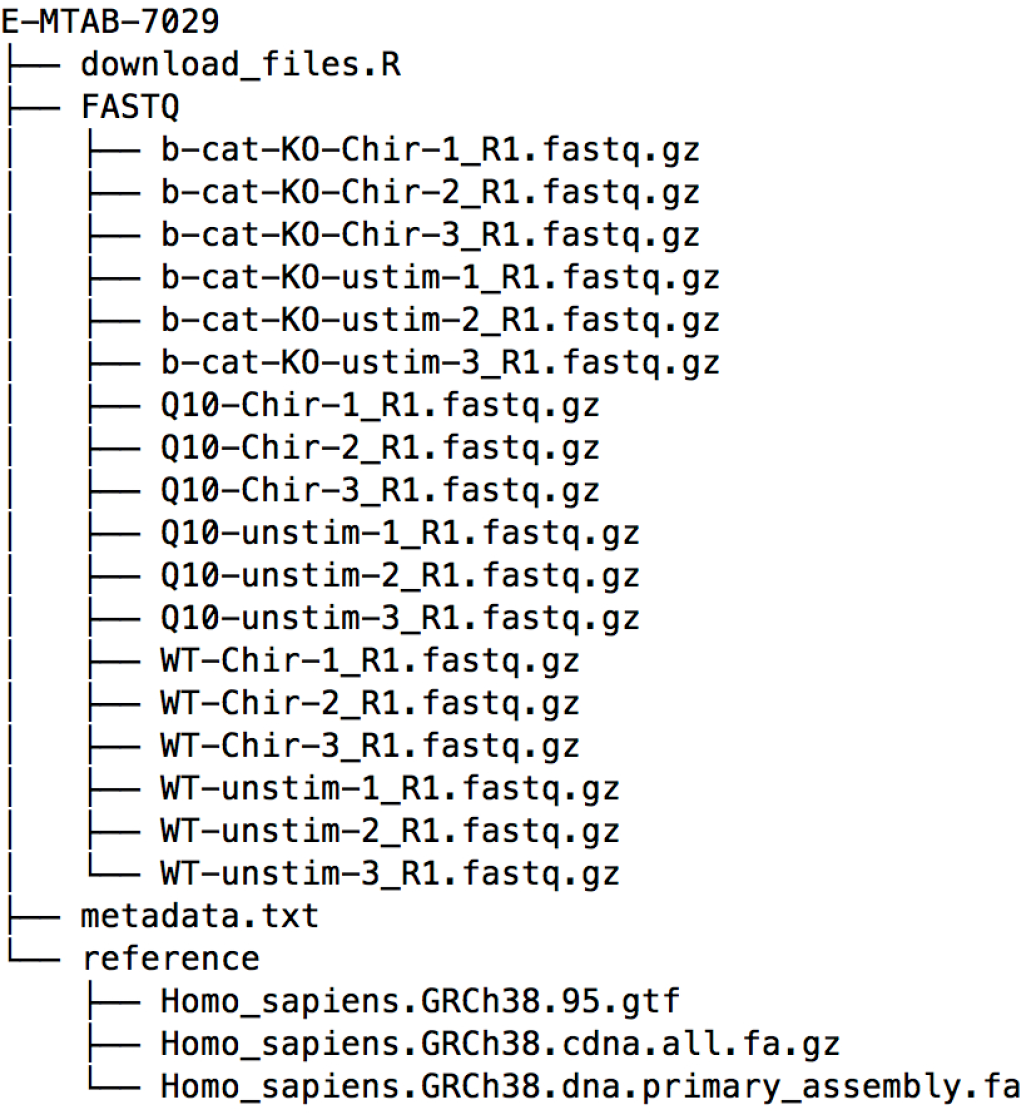
The suggested structure for the set of files that need to be organized to run ARMOR on a new dataset. The structure can deviate from this somewhat, since the location of the files can be specified in the corresponding config.yaml file.

The set of files (not including the large data and reference files, which would be downloaded using the `download_files.R`) used in this setup can be found on the chiron_realdataworkflow branch of the ARMOR repository: https://github.com/csoneson/ARMOR/tree/chiron_realdataworkflow.

After downloading the data, generating the metadata.txt file and editing the config.yaml file, the full workflow was run with the command:

~~~
snakemake --use-conda --cores 6 \
--configfile config_E-MTAB-7029.yaml
~~~

and after some hours of computation, the specified output directory was populated as shown in Figure 4. Using the template run_iSEE.R (from the chiron_realdataworkflow branch of the ARMOR GitHub repository) and the output file shiny_sce.rds from running ARMOR, an R/shiny web application can be initiated, with various panels to allow the user to interactively explore the data and results (Figure 5).

## Acknowledgments

M.D.R. acknowledges support from the University Research Priority Program Evolution in Action at the University of Zurich, the Swiss National Science Foundation (grants 310030_175841, CRSII5_177208) and the Chan Zuckerberg Initiative (grant 2018-182828).

---

**Figure 4:**
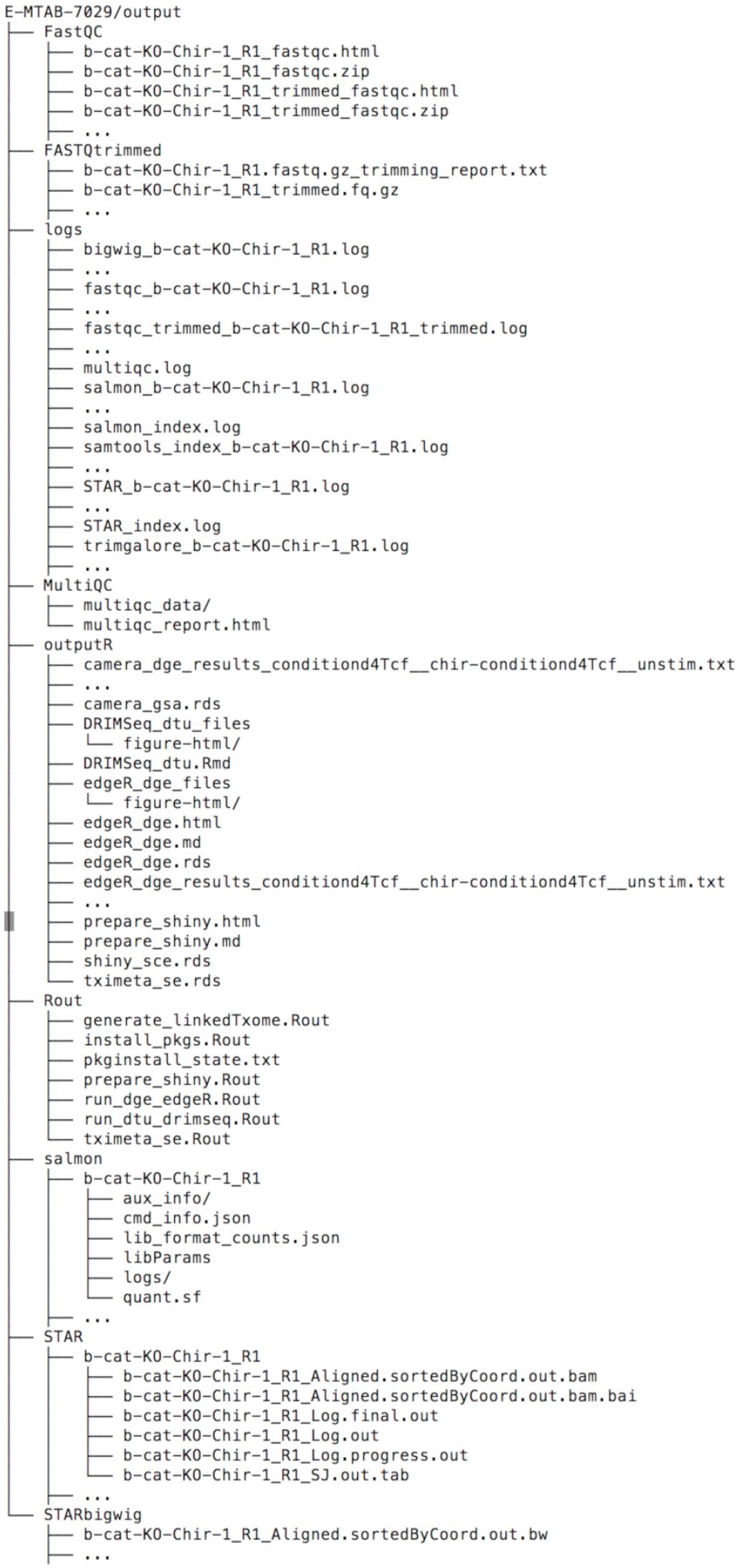
The set of output files from the workflow. This includes log files for every step and all the standard outputs of all the tools, such as R objects and scripts, BAM files, bigWig files and quantification tables. Note that the outputs for only one RNA-seq sample are shown; … represents the set of output files for the remaining samples or contrasts and directories ending in / contain extraneous files and are collapsed here.

**Figure 5:**
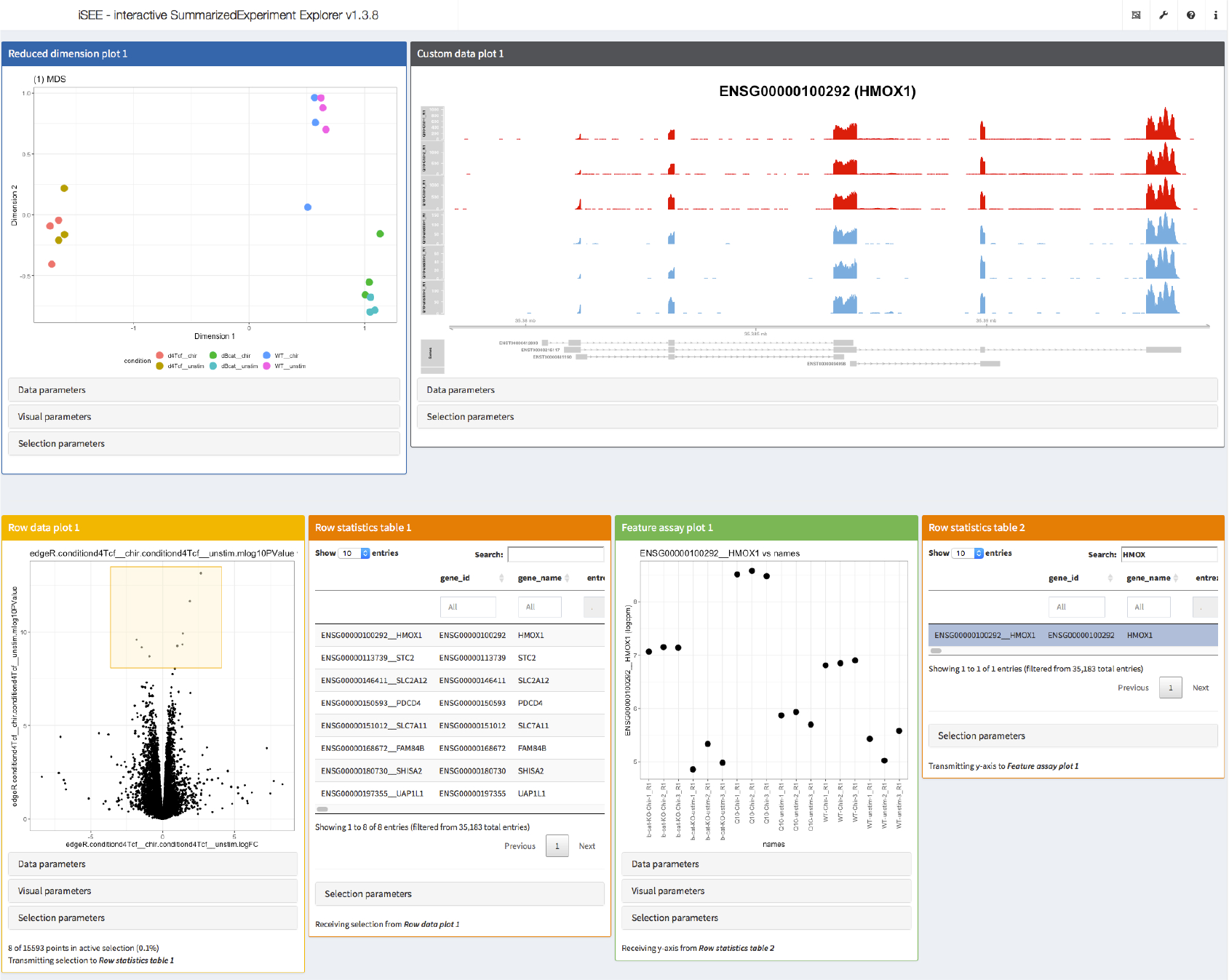
Screenshot of visualization of data and results from the real data walk-through using the iSEE R/Bioconductor package. The interactive application was configured to display an MDS plot colored by the sample condition (top left), a custom panel showing the observed read coverage of a selected gene (top right), a volcano plot for a specified contrast (bottom left, the selected genes are shown in the adjacent table) and an overview of the log-CPM expression values for each sample, for a gene selected in a second table (bottom right).

